# Seven blood biomarkers are associated with TGF-β and VHL-HIF signaling in patients with clear cell renal cell carcinoma

**DOI:** 10.1101/2025.02.07.636864

**Authors:** Pramod Mallikarjuna, Cemal Erdem, Harald Hedman, Raviprakash T. Sitaram, Ruben Ilundain Beorlegui, Karthik Aripaka, Anders Larsson, Börje Ljungberg, Masood Kamali-Moghaddam, Maréne Landström

## Abstract

Clear cell renal cell carcinoma (ccRCC) is an aggressive kidney cancer subtype frequently associated with poor prognosis. Most ccRCC cases are asymptomatic in early stages and symptomatic mostly in advanced stages. Furthermore, the heterogeneity of ccRCC presents a challenge to design new treatments. In this study, using proximity extension assay (PEA), we analyzed blood samples from 134 patients with ccRCC and from 111 age- and gender-matched healthy donors. We identified a panel of seven proteins (ANXA1, ESM1, FGFBP1, MDK, METAP2, SDC1, and TFPI2) that are associated with clinicopathological parameters and patient survival. These biomarkers can differentiate patients with ccRCC from the control individuals with high diagnostic sensitivity and specificity. Moreover, by studying protein expression in solid tumors from the same ccRCC patients, we revealed associations between the panel biomarkers and proteins in the TGF-β and VHL-HIF signaling pathways. We found that most tumor promoting biomarkers were positively associated with TGF-β signaling and HIF-2α, and negatively associated with pVHL and HIF-1α. We also found that most tumor suppressing biomarkers were positively associated with pVHL and HIF-1α and negatively associated with TGF-β signaling and HIF-2α. For ccRCC patients, the blood protein biomarkers that were connected to poor prognosis and TGF-β/HIF-2α signaling, as identified in this study, are potentially important assets in personalized medicine.

## Introduction

Renal cell carcinoma (RCC), accounts for 2.2% of all cancers diagnosed worldwide, and the mortality due to this disease constitutes 1.8% of all cancer-related deaths^1^. RCC was previously often presented by hematuria, flank pain, and palpable abdominal mass, which are considered the classic triad of symptoms for all RCC subtypes including clear cell renal cell carcinoma (ccRCC) - the most common RCC type^2^. However, only a small percentage of patients present these symptoms today, and RCC is therefore often diagnosed incidentally due to an investigation for other clinical conditions^3^. One third of the localized diseases develop metastasis during oncologic follow-up, and half of the patients with post-nephrectomy recurrence were associated with cancer-specific death^4^. ccRCC constitutes about 70% of all RCCs, sporadically associated with familial syndromes such as von Hippel-Lindau (VHL) disease^5,6^. Although the benefits of early detection of cancer are immeasurable and some molecular biomarkers have emerged as new tools in precision medicine^7^, specific markers for ccRCC are still quite scarce.

Recently, it has been accentuated that immunohistochemistry for CA9, a downstream target gene in the hypoxia pathway, is the best available marker for ccRCC, performed in tissue biopsies or tissues from the tumor^8–10^. On the other hand, the liquid biopsies offer the advantage of non-invasive sampling and the possibility of longitudinal monitoring for patients with ccRCC^11^. However, currently there are no validated liquid biopsy assays available for RCC. In this study, we explore the potency of proximity extension assay (PEA) for the detection of marker proteins in blood samples from patients with ccRCC. PEA offers the advantage of sensitive detection of proteins in plasma from minute amounts of blood^12^. We used the Olink Target 96 Oncology II PEA-panel^13^ to measure protein levels in serum from patients diagnosed with ccRCC. As control, we used blood serum from age- and sex-matched healthy individuals. The PEA datasets were analyzed using machine learning algorithms to identify a minimal ccRCC signature protein panel that can be used for identification of patients at risk of developing ccRCC.

Previously, we have reported that the transforming growth factor-β (TGF-β) and VHL signaling components interact and affect each other in RCC^14^. TGF-β is a key regulatory cytokine required for proper embryogenesis, differentiation, and tissue homeostasis as well as maintenance of a normal immune response in healthy individuals, and its dysregulation leads to cancer and autoimmune diseases^15^. TGF-β binding to its tetrameric receptor complex at the plasma membrane leads to activation of signaling cascades through Smad-dependent (canonical) or Smad-independent (noncanonical) pathways^15,16^. Both the canonical and noncanonical components of TGF-β signaling contributed to ccRCC progression^17–19^. In the noncanonical TGF-β signaling pathway, we have previously reported that the TGF-β Type 1 receptor undergoes proteolytic cleavage in prostate cancer and ccRCC cells, resulting in an intracellular domain which enters the nucleus and promotes an invasive program^17,20,21^. VHL, a tumor suppressor often mutated in ccRCC^22^, is a E3 ubiquitin ligase that targets and marks the tumor-promoting hypoxia-inducible factors HIF-1α and HIF-2α for degradation by the proteasome in an oxygen-dependent manner^23^. We have shown that TGF-β contributes to the aggressiveness of ccRCC dependent on the VHL mutation status^18^. The TGF-β targets plasminogen activator inhibitor-1 (PAI-1) and SNAIL1 are also associated with ccRCC progression^17,19^. In the second part of the current study, we used immunoblotting of tumor cell lysates to explore the connections of the signature proteins with the TGF-β and hypoxia/VHL signaling pathways^17–19^.

In this study, we identified a minimal subset of blood serum biomarkers to classify patients with ccRCC or those who have a high risk to develop metastatic ccRCC. The second aim of the study was to investigate if the biomarkers are connected to the TGF-β and hypoxia pathways in ccRCC. Our current study complements our previous findings, and we present here several novel connections between blood biomarkers and TGF-β/hypoxia signaling components within ccRCC tumors.

## Results

### The ccRCC patient cohort and measurement of blood plasma protein levels

The experimental setup and sampling procedure of patients and control participants is described in Figure 1A. The tumor size of each patient (N=142) was determined by computed tomography (CT). The mean tumor diameter in the patient group was 70 mm (range 12-190 mm). The tumor stage distribution was determined according to the 2009 TNM classification system^24^ and was as follows: 58 patients in TNM stage I (40.8%), 23 patients in stage II (16.2%), 28 patients in stage III (19.7%), and 33 patients in stage IV (23.2%). The Fuhrman *et al*.^25^ grading system was used to grade the tumors, with the following grade distribution: 18 (12.7%) tumors were grade 1, 54 (38.0%) grade 2, 46 (32.4%) grade 3, and 24 (16.9%) grade 4. Patient follow-up performed using a scheduled program was employed for survival analysis. As of October 2022, 36 (25.4%) patients with ccRCC were alive without any indication of disease, 4 (2.8%) were alive with disease, 58 (40.8%) had died of ccRCC, and 44 (31.0%) had died of other causes.

**Fig. 1.**
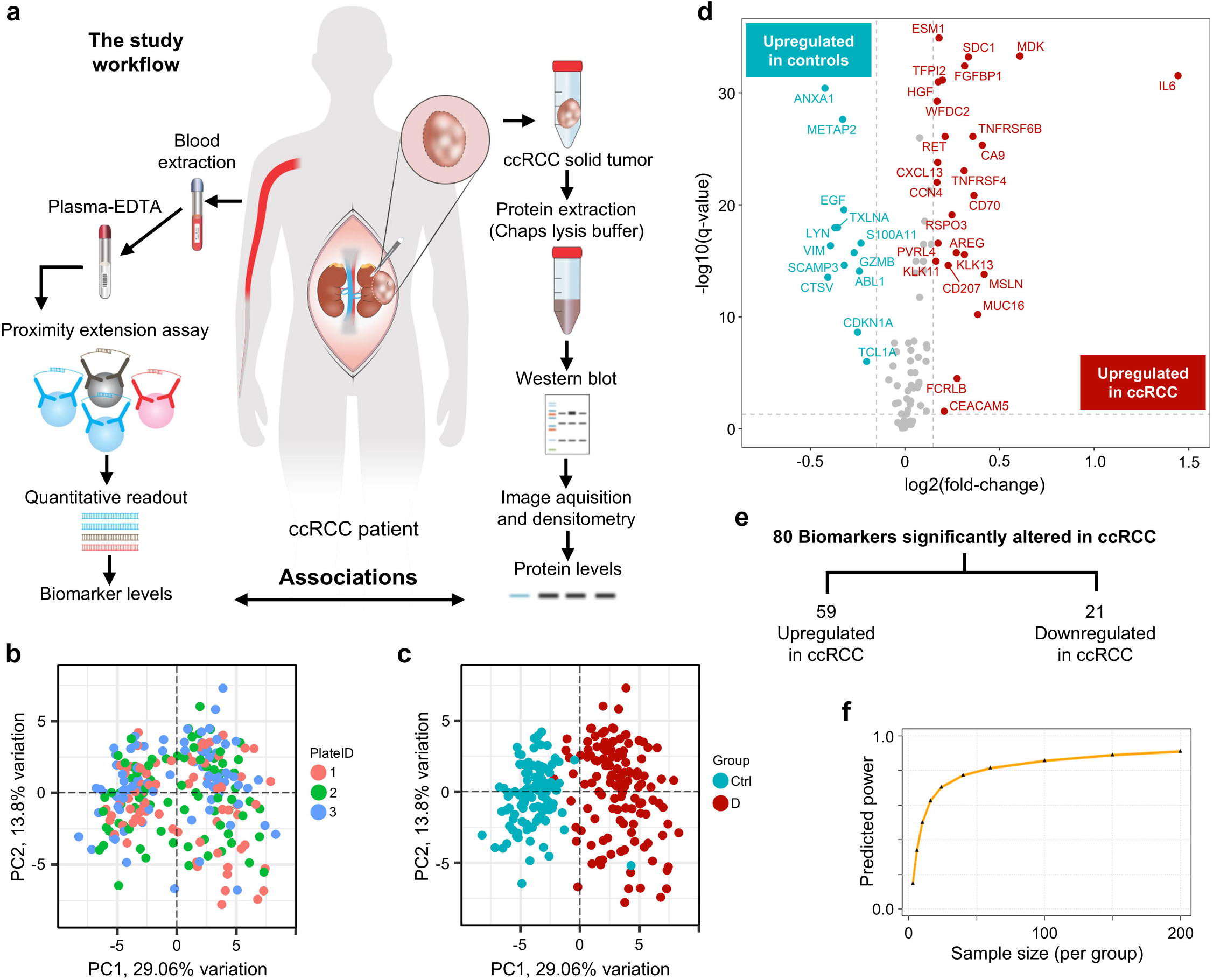
The serum biomarkers are differentially expressed in ccRCC patients compared to healthy controls. **a** The schematic illustration of the two parts of the study, which focuses on identifying novel biomarkers in the blood of patients with ccRCC. Additional analysis has been performed to investigate the associations between TGF-β and hypoxia pathway proteins from solid ccRCC tumors of the same patient cohort. **b** Principal component analysis (PCA) of NPX data from three assay plates (1, 2, or 3) of blood samples from ccRCC patients and healthy controls shows lack of distinct cluster groups among the plates, suggesting negligible technical bias. **c** PCA of NPX data from ccRCC patients and healthy controls shows a clear separation between the ccRCC samples (D, red circles) and healthy controls (Ctrl, blue circles). **d** Volcano plot summarizing the abundance of biomarkers significantly altered in ccRCC samples compared to healthy controls. Two-sided, unpaired Wilcoxon Rank Sum test, p-values are FDR corrected (Supplementary Data 3). Only biomarkers with log2 fold-change greater than 0.15 or less than -0.15 with adjusted p-value<0.05 are highlighted and labelled. **e** The numbers of biomarkers differentially expressed in ccRCC patients and healthy controls. Significance at FDR corrected p-values<0.05. **f** Predicted power curve showing the statistical power imparted by the number of samples used in this study. With 111 healthy controls and 134 patients, the predictive power is around 0.85-0.9.

Blood plasma protein levels of patients (N=134) and healthy controls (N=111) were quantified using proximity extension assay (PEA) of Olink Target 96 Oncology II panel (Olink Proteomics, Uppsala, Sweden). Eight of the initial 142 patients and 10 of the initial 121 healthy samples were not quantifiable by the assay and thus excluded from the dataset for further analyses. A total of 92 oncology-related protein biomarker candidate levels were measured across all samples and the Olink-defined Normalized Protein eXpression (NPX) units were used (log2 scale) (Supplementary Data 1).

### Evaluation of technical bias and covariates

First, we filtered the Olink dataset and excluded proteins with NPX levels below the limit of detection (LOD) in more than 60% of samples. Only the FADD protein measurements met this criterion, removed from the input data, and the remaining 91 proteins were retained for further analyses.

Next, linear regression against clinically relevant features identified 12 proteins that were affected by patient age (p < 0.05): CCN4, CD27, CXL17, CYR61, EPHA2, IGF1R, ITGB5, RET, TGFR2, TNFRSF19, TNFSF13, and WFDC2. No significant effect was observed for the patient’s gender. A final age-adjusted dataset was produced by adjusting the values to account for the effect of age on protein expression (Supplementary Data 2 and Supplementary Figures 1-12).

Then, we explored inter-assay variation between the three PEA plates. We used principal component analysis (PCA) and saw a lack of distinct cluster groups, meaning no technical bias detected (Figure 1B). A subsequent PCA was performed to evaluate the potential to distinguish between the control and disease groups. A very clear separation was observed between the tumor samples and the controls (Figure 1C).

### Identification of differentially expressed proteins and association of the biomarkers with clinicopathological parameters

Of the 91 biomarkers examined, the expression levels of 80 proteins were significantly altered in ccRCC patients compared to controls (Figure 1D and Supplementary Data 3). The abundance of proteins significantly altered in tumor samples compared to controls had 59 upregulated and 21 downregulated proteins (Figures 1D and E).

The biomarkers significantly associated (p < 0.05) with higher grade, advanced stage, and larger tumors are categorized as tumor promoters, whereas biomarkers associated with lower grade, earlier stage, and smaller tumors are categorized as tumor suppressors. A few biomarkers were not significantly associated with any of the clinicopathological parameters and were therefore neither promoters nor suppressors. 35 proteins are categorized as tumor promoters, 12 as tumor suppressors, and 33 as neither promoters nor suppressors (Figures 2A and B). To further understand the role of the biomarkers, we next examined whether their differential expression in the healthy and ccRCC cohorts aligned with their status as tumor promoters, tumor suppressors, or neither (Figure 2C). The biomarkers are either positively or negatively associated with cancer-specific survival. Most of the tumor promoters are associated with poor survival and upregulated in patients compared to healthy controls (Figure 2C).

**Fig. 2.**
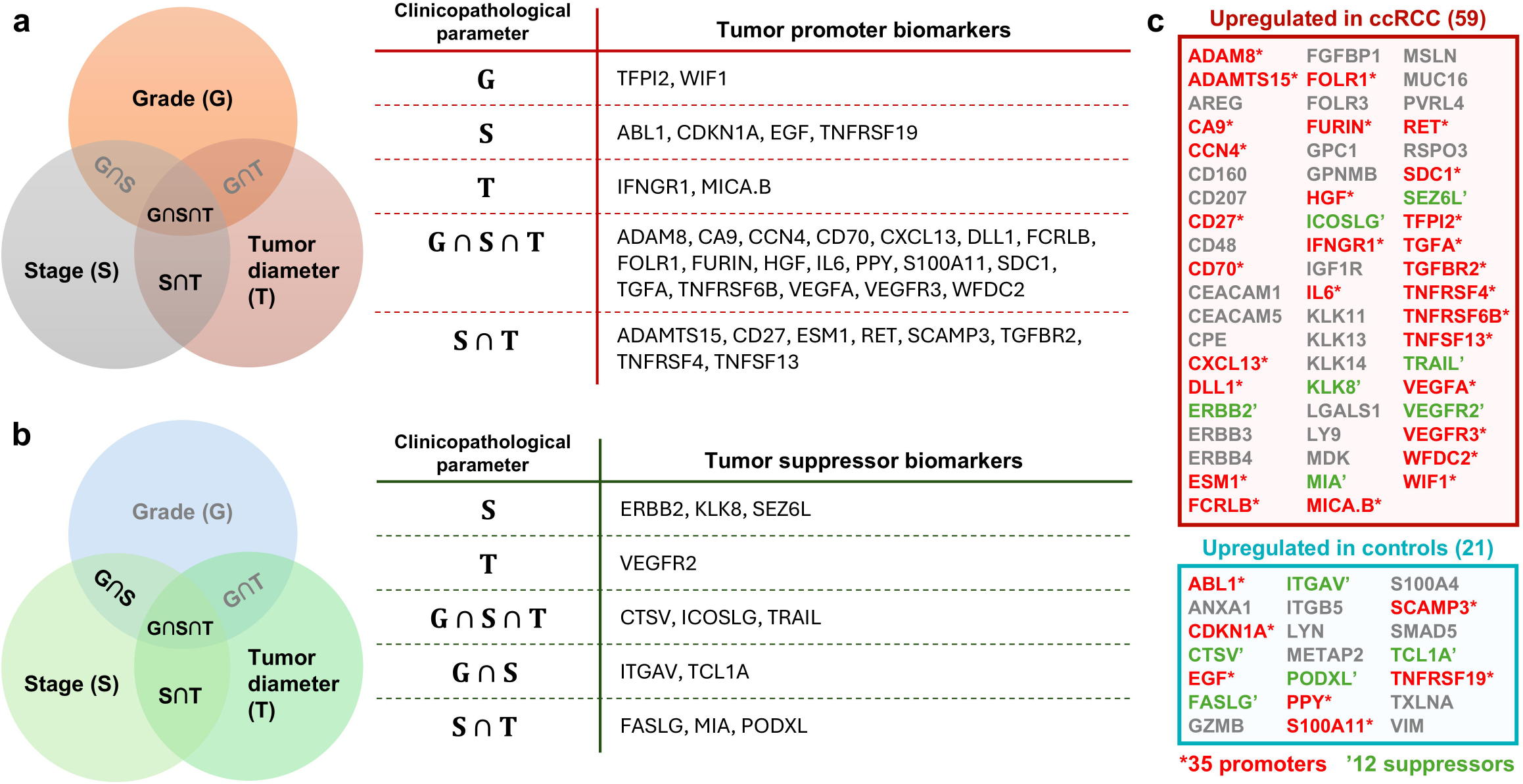
Associations of the biomarkers with tumor grade, stage, and size cluster them into different groups. **a** Venn diagram representing the association of biomarkers with higher grade, advanced stage, and larger tumors. The biomarkers listed in this figure are associated with tumor promotion. Significance at FDR corrected p-values<0.05, Spearman correlation. **b** Venn diagrams representing the association of biomarkers with lower grade, early stage, and smaller tumors. The biomarkers listed in this figure are associated with tumor suppression. Significance at FDR corrected p-values<0.05, Spearman correlation. **c** All 80 blood biomarkers significantly altered in this study are classified into tumor promoters, tumor suppressors, and neither tumor promoters nor tumor suppressors, based on association/no-association of the biomarkers with grade, stage, and tumor diameter (**a** and **b**).

### Identification of a minimum ccRCC signature panel

To evaluate the potential of the proteins in distinguishing tumor samples from control samples, we used the top 50 most significantly altered proteins to train a random forest (RF) model. The importance of each protein biomarker for classifying the data was then inspected (Figure 3A), where cross-validation model performances were close to 100% (median=100%, mean=99.40%, Figure 3B, and Supplementary Data 4-5).

**Fig. 3.**
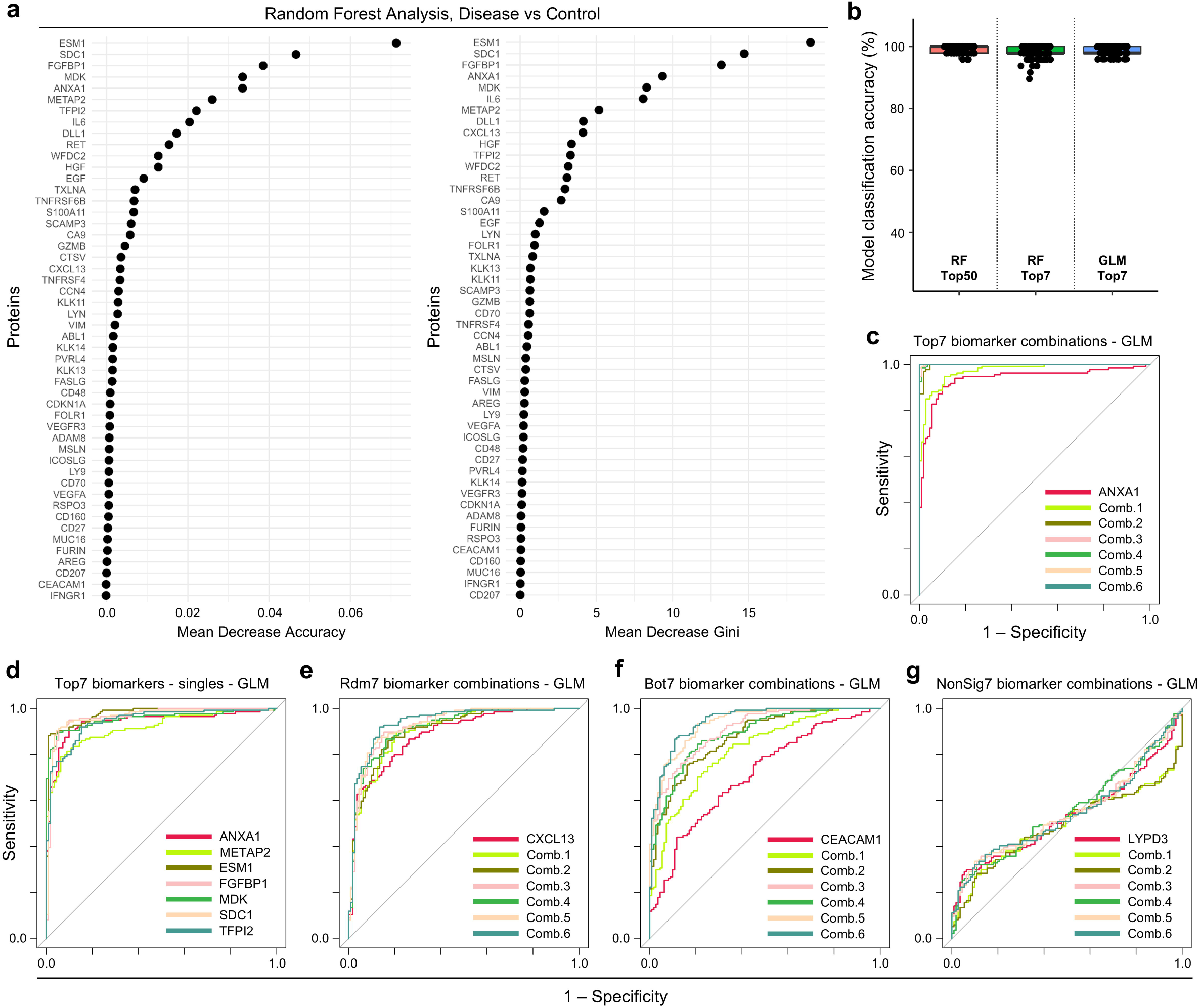
Machine learning shows the importance of individual biomarkers in correctly classifying the patients and controls. **a** The left panel shows the mean of the model accuracy decrease and the right panel shows the decrease in Gini coefficients, per biomarker tested in the random forest model (regression model, age-adjusted data, top 50 significantly altered proteins, number of trees=100). **b** The validation set accuracy for different machine learning models in classifying patients. Using the top 50 differentially expressed biomarkers and random forest classification, the model accuracy is close to 100% for all 100 models tested (red bar). Using only the top seven biomarkers are also close in accuracy with some models performing around 95% (green bar). Using gaussian linear models (GLM) with the top seven biomarkers also perform with accuracy closer to 100% (blue bar). **c** The ROC curve for correctly classifying ccRCC patients and healthy controls. The red curve shows the model performance when only ANXA1 data is used. Each new curve shows the model performance with an additional biomarker, up to all seven. The model with all seven is a perfect classifier (AUC = 1). **d** The models with only individual biomarkers, of the top seven, are not better than the combination in **c**. **e** The combination model performances of seven randomly selected biomarkers from the top 50 significantly altered biomarker list does not perform at 100% accuracy. **f** As expected, the least significantly altered proteins from the top 50 list perform worse in correctly identifying the patients or controls. **g** Finally, the combination models of the seven biomarkers that are not significantly altered between patients and healthy controls cannot correctly classify ccRCC patients.

We next sought to identify the smallest subgroup of serum biomarkers that could be used to produce an accurate and precise diagnostic model for ccRCC. Based on the mean accuracy decrease metric (Figure 3A), we systematically included increasing number of proteins in a new RF model and saw that the top seven are enough to perfectly classify patients and healthy controls (AUC=1, Supplementary Data 6). The ROC curves of combinations of proteins were compared, where the proteins with the highest absolute regression coefficients were added together iteratively until no further significant improvement in AUC value was obtained with the addition of more proteins. The cross-validation accuracies of the top seven biomarker models were also close 100% (median=97.62%, mean=98.08%, Figure 3B) and the performance of the combined signature was greater than that of any of the seven individual proteins. The final list of proteins is: ANXA1, ESM1, FGFBP1, MDK, METAP2, SDC1, and TFPI2.

### The performance of the seven-gene ccRCC signature panel

To further test the performance of the identified seven gene signature, we trained an elastic-net penalized logistic regression (ENLR) model in R package glmnet^26^. The ENLR model combines lasso and ridge regularization penalties to simultaneously perform regularization and variable selection. Similarly to the RF model testing, the combination of seven biomarkers resulted in a perfect classifier (Figure 3C) and was better than using any of the seven individually (Figure 3D). Furthermore, the selection of a random set of seven proteins and their combinations were also not as significant as the chosen top seven (Figure 3C vs 3E). The accuracy of combining the bottom seven of the top 50 significantly altered protein levels were even less accurate (Figure 3F) but better than combining the least altered seven proteins (Figure 3G). Comparing 100 cross-validation model performances of the top seven (Figure 3B) vs bottom seven, the non-significant seven, and 10 randomly selected seven proteins (Supplementary Figure 13), we showed that the chosen seven protein signature panel is the most accurate.

The performance differences between the sets of random seven protein panels were partly based on inclusion of one or more of the top seven genes. The lists of genes selected and the validation data with predicted classes are given in Supplementary Data 7-10. Finally, a model of seven non-significantly altered proteins in classifying patients showed that it is the not better than random (Figure 3G) and confirmed our approach of selecting top highly and significantly altered proteins as the signature panel.

Kaplan-Meier survival plots were plotted to further investigate the association of these biomarkers with cancer-specific survival, higher expression of five of these seven biomarkers were significantly associated with poor cancer-specific survival (Figure 4). We observed that ESM1, METAP2, SDC1, and TFPI2 are associated with poor cancer-specific survival and FGFBP1 with better survival, whereas ANXA1 and MDK are not associated with survival. The identification of ESM1, METAP2, SDC1 and TFPI2 as potential blood biomarkers for poor prognosis for patients with ccRCC gives support to follow up our data in future longitudinal studies.

**Fig. 4.**
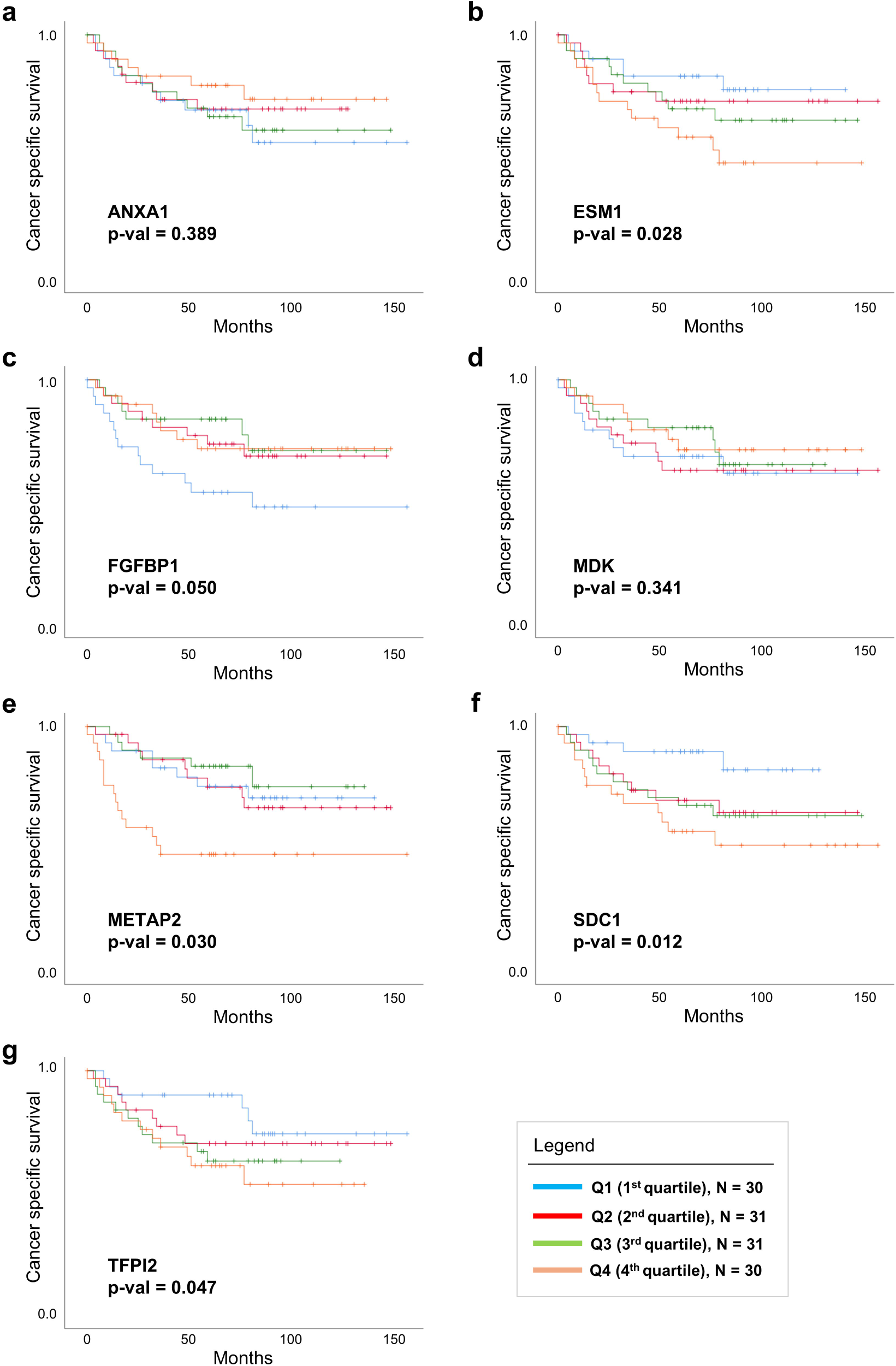
Kaplan-Meier survival plots for the top seven biomarkers show cancer specific survival. **a** ANXA1. **b** ESM1. **c** FGFBP1. **d** MDK. **e** METAP2. **f** SDC1. **g** TFPI2. **(a-g)** The follow-up patient cohort (N=122) is divided into four quartiles based on biomarker expression levels and cancer specific survival rates between the groups are compared: Q1vs Q2 vs Q3 vs Q4, Q1 vs Q2+Q3+Q4, Q1+Q2 vs Q3+Q4, and Q1+Q2+Q3 Vs Q4. Significance reported at p-values<0.05.

### Associations of TGF-β signaling components with plasma protein biomarkers

We have previously reported that the TGF-β signaling and the hypoxia pathways are interconnected and involved in ccRCC^17–19^. Here we explore the connections between these two pathways and the proteins identified using the PEA analysis (Figure 1A). The immunoblot experiment revealed that full length TGFBR1 (TGFBR1-FL) is positively correlated with the tumor promoters VEGFA, EGF, IL6, NT5E, ADAMTS15, and FCRLB and is negatively correlated with tumor suppressors ERBB2, SEZ6L, ICOSLG, GPNMB, and FGFBP1 (Figure 5B and Supplementary Data 11-12).

**Fig. 5.**
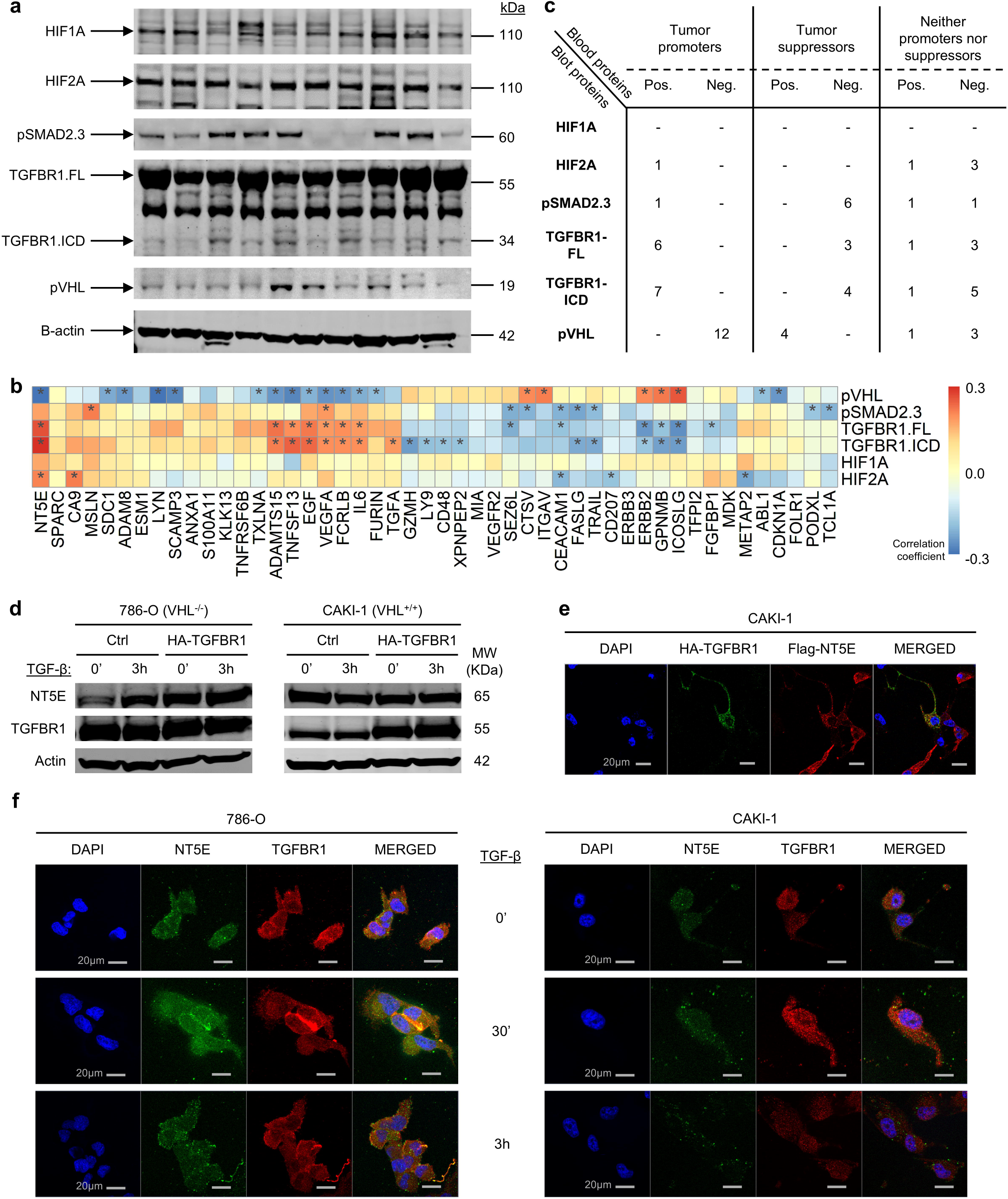
The serum biomarker levels are significantly correlated with TGF-β and HIF-α/pVHL pathway proteins in solid tumors. **a** Representative western blot image showing the protein bands of HIF1A, HIF2A, pSMAD2/3, TGFBR1-FL, TGFBR1-ICD, pVHL, and β-actin, in total protein extracted from ccRCC solid tumor excisions. **b** Correlation matrix representing correlations between protein components of TGF-β and HIF-α/pVHL pathway proteins with serum biomarkers. * denotes significance at p-values<0.05, Spearman correlation. **c** The table summarizes the numbers of biomarkers significantly associated with the TGF-β and HIF-α/pVHL pathway proteins, based on the biomarker classifications (Fig. 2). **d** Transient overexpression of TGFBR1 or stimulation with TGF-β induces NT5E in VHL-negative 786-O cells but not in VHL-positive CAKI-1 cells. **e** Transiently overexpressed TGFBR1 (HA-TGFBR1, green) and NT5E (red) proteins are colocalized in CAKI-1 cell membrane and cytoplasm (merged, yellow). DAPI (blue) stain represents nuclear regions. The scalebar in grey shows 20μm. **f** Endogenous TGFBR1 (red) and NT5E (green) are colocalized (merged, yellow) in the two ccRCC cell lines treated with TGF-β, where the two proteins are likely to form protein-protein complexes. DAPI (blue) stain represents nuclear regions. The scalebar in grey shows 20μm. Rows correspond to TGF-β stimulation of 0min, 30min, and 3hours from top to bottom, respectively.

The immunoblotting measurements for only the intracellular domain of TGFBR1 (TGFBR1-ICD originally described in Mu et al., 2011^20^ and Gudey et al., 2014^21^) are correlated positively with the tumor promoters VEGFA, EGF, IL6, NT5E, ADAMTS15, FCRLB, and TGF-α and negatively with the tumor suppressors ERBB2, ICOSLG, GPNMB, FASLG, and TRAIL (Figure 5B and Supplementary Data 11-12).

The pSMAD2/3 proteins are correlated positively with the tumor promoters VEGFA and MSLN and negatively with the tumor suppressors SEZ6L, FASLG, TRAIL, CTSV, PODXL, and TCL1A (Figure 5B and Supplementary Data 11-12). Other correlations of TGFBR1-FL, TGFBR1-ICD, and pSMAD2/3 with biomarkers that are neither tumor promoters nor tumor suppressors are given in Supplementary Data 11-12.

### Associations of hypoxia signaling components with plasma protein biomarkers

The immunoblot experiment with the hypoxia pathway proteins revealed that HIF-1α (HIF1A) was not significantly associated with any of the biomarkers but still showed negative correlations with tumor promoters ADAMTS15 and TNFSF13 (Figure 5B and Supplementary Data 11-12). HIF-2α (HIF2A) was positively associated with the following tumor promoters: NT5E and CA9 (Figure 5B and Supplementary Data 11-12). pVHL was positively associated with the tumor suppressors CTSV, ICOSLG, ERBB2, GPNMB, and ITGAV and negatively with the tumor promoters ADAMTS15, SDC1, TNFSF13, NT5E, TXLNA, VEGFA, EGF, IL6, FCRLB, FURIN, SCAMP3, ADAM8, CDKN1A, and ABL1 (Figure 5B and Supplementary Data 11-12). Other correlations of HIF-1α, HIF-2α, and pVHL with biomarkers that are neither tumor promoters nor tumor suppressors are given in Supplementary Data 11-12.

### Summary of associations between TGF-**β** signaling, HIFs, and plasma proteins of ccRCC patients

There are significant associations between TGF-β and hypoxia pathway proteins with biomarkers (Figure 5C). Furthermore, all components of TGF-β signaling and HIF-2α are positively associated with tumor promoters and negatively associated with tumor suppressors; in contrast, HIF-1α and pVHL are positively associated with tumor suppressors and negatively associated with tumor promoters.

### Influence of TGF-**β** and levels of TGF-**β** Type I receptor on expression of NT5E in ccRCC cells

Since we observed a positive correlation between expression levels of TGFBR1 in solid tumor tissues and NT5E as a biomarker in blood from patients with ccRCC (Figure 5B), we examined if transient overexpression of TGFBR1 and stimulation with TGF-β could influence the expression of NT5E using two ccRCC cell lines, by using immunoblotting (Figure 5D). We observed that both TGF-β stimulation of 786-O cells, or transient overexpression of TGFBR1, increased the expression of NT5E. In contrast, the positive effect of TGF-β stimulation on NT5E levels was not so prominent in CAKI-1 cells, which express VHL. Moreover, we also investigated a possible association between transiently overexpressed TGFBR1 and NT5E in CAKI-1, by using co-immunofluorescence. As shown in Figure 5E, we observed that the two proteins were found to be colocalized in cell membrane and in the cytoplasm. We examined also if endogenous TGFBR1 and NT5E could be found to be colocalized by co-immunofluorescence in the two ccRCC cell lines treated with TGF-β (Figure 5F). From these experiments we conclude that TGF-β stimulation of ccRCC cells could have a positive effect on NT5E expression and that TGFBR1 and NT5E are forming a protein-protein complex in the cell membrane and cytoplasm.

## Discussion

This study had the outset to identify liquid biomarkers that can be used for the stratification of patients suffering from early or late stages of ccRCC. Protein expression data was obtained using a blood biobank collected in northern Sweden. In short, blood protein levels were measured by PLA using the Olink Oncology-II panel and analyzed by machine learning models to identify new biomarkers as collective indicators of ccRCC. The Olink panel proteins were clustered into three groups; proteins associated with ccRCC tumor promotion (35) or suppression (12) and proteins associated with neither tumor promotion nor suppression (33). Interestingly, there was no overlap between proteins associated with early stage/low grade tumors and proteins associated advanced grade tumors at a later stage. Proteins associated with poor cancer-specific survival were tumor promoters, and proteins associated with better cancer-specific survival were identified as tumor suppressors (Figure 2). The statistical analyses identified the following seven signature proteins that all were associated with ccRCC: ANXA1, ESM1, FGFBP1, METAP2, MDK, SDC1, and TFPI2.

FGFBP1 (fibroblast growth factor binding protein 1) is expressed in epithelial cells in skin, stomach, eye, ileum, and colon and it is upregulated in various types of cancers^27^. FGFBP1 is a secreted chaperone that enhances the activity of locally secreted fibroblast growth factors by presenting these to nearby cells expressing FGF receptors. In our data, serum levels of FGFBP1 were increased in ccRCC patients (Figure 1D), suggesting that FGFBP1 can function as a promoter of aggressive ccRCC which warrants further studies. Intriguingly, it has been shown that TGF-β inhibits FGFBP1 expression in skeletal muscle^28^. We conclude that FGFBP1, as a secreted protein, is a promising blood biomarker for ccRCC in an early stage and might be useful for stratification of patients that will require less aggressive interventions.

We found that high levels of ESM1 (Endothelial cell-specific molecule 1) or endocan, showed a strong correlation to poor survival. ESM1 is a key factor in vascular development, neogenesis, and angiogenesis, and is secreted by vascular endothelial cells, playing an important role in endothelium-dependent pathological disorders and inflammatory reactions^29^. In our study, ESM1 was identified as a tumor promoter, with an increased expression in plasma from ccRCC patients. These results are supported by other studies, reporting that ESM1 is dramatically overexpressed in many cancer types including non-small-cell lung cancer, clear cell renal cell carcinoma, and ovarian cancer^30^. Supporting our findings here, it was shown previously that vascular ESM1 is markedly overexpressed in ccRCC^31^, and the study by Kim et al.^32^ reported significantly higher ESM1 levels in ccRCC patients with large tumors and higher stages. It was concluded that ESM1 can serve as a serologic biomarker for diagnosing and monitoring RCC, particularly in patients younger than 50 years^32^.

METAP2 (Methionine aminopeptidase 2) showed a reduced expression in ccRCC patients and was found to be associated with poor survival of ccRCC patients in our study (Figures 1D and 4E). METAP2 catalyzes the removal of N-terminal methionine from nascent proteins, and in most cellular proteins this removal is required for proper protein function^33^. METAP2 has been a drug target for over three decades, with multiple pharmaceutical companies competing in the quest for potent METAP2 inhibitor for the treatment of cancer and more recently obesity and autoimmunity^34^. To our knowledge this is the first study that identifies METAP2 as a serum biomarker for ccRCC.

Another potential liquid biomarker for ccRCC patients identified in this study is SDC1 (Syndecan-1) a transmembrane heparan sulfate proteoglycan protein that mediates a cellular response to several paracrine signals^35^. SDC1 has been reported as a marker for various cancers^36^. Here we identified SDC1 as a biomarker for aggressive ccRCC (Figures 2A and 4F). This finding contrasts with the study of Niedworok et al.^37^, that found no correlation between serum or tissue levels of SDC1 and the prognosis for patients diagnosed with RCC.

Similar to FGFBP1 and METAP2, ANXA1 (Annexin A1) and MDK (Midkine) were also not identified as a promoter or repressor of tumors in our analyses. Although ANXA1 showed markedly reduced expression in tumors of ccRCC patients compared to controls (Figure 1D), its expression was not significantly associated with clinicopathological parameters or cancer-specific survival (Figures 2C and 4A). ANXA1 is the founding member of the annexin superfamily of calcium-dependent phospholipid-binding proteins. ANXA1 expression is regulated by glucocorticoids and inhibits the action of cytosolic phospholipase A2 (PLA2) resulting in a reduced synthesis of eicosanoids^38^. In contrast to our findings, elevated levels of ANXA1 have been detected in lung cancer, colorectal cancer, hepatocellular carcinoma, pancreatic cancer and in melanomas^38,39^. The upregulation of ANXA1 is related to malignancy of some cancer forms and ANXA1 was detected in blood vessels of the human prostate, liver, breast, and lung tumors, but not matched normal tissues^39^. Our finding of lowered ANXA1 levels in ccRCC patients is in contradiction to the report from Yamanoi^39^, in which high expression of ANXA1 correlated with malignant potential in RCC patients. Also, for other cancer types, high expression of ANXA1 is positively correlated with disease severity and an increasing tumor stage. However, the effect of ANXA1 on tumor proliferation is not fully understood, and ANXA1 can function both as a suppressor and promotor of cancer^38^. ANXA1 has also gained interest as a therapeutic target, where in a murine model for melanoma, lowered levels of the ANXA1 reduced the invasive potential of cancer cells^38^. In short, our findings linking reduced levels of ANXA1 in serum to ccRCC warrants more detailed studies.

In other malignancies, MDK was present at a higher level in the plasma of the ccRCC patients but was not associated with any clinicopathological parameters or patient survival. MDK is a secreted growth factor with a high homology to Pleiotropin (PTN), both sharing common receptors^40^. MDK has a temporary high expression during embryogenesis, particularly in the kidneys, which is reduced to low levels in adult tissues^41^. MDK has been reported as a marker for cancer, and its expression is detected in precancerous stages of human colorectal and prostate carcinomas. At advanced stages, most carcinoma specimens express MDK at a high level, and in more than 60% of human adult carcinomas, an elevated serum MDK level was detected^40^. MDK is also expressed in inflammatory cells and is released by endothelial cells under hypoxic conditions^41^. These reports support the identification of MDK as biomarker for ccRCC.

The last biomarker of the identified signature panel is TFPI2 (Tissue Factor Pathway Inhibitor 2), which is a serine protease inhibitor. It is involved in various physiological processes including coagulation and fibrinolysis. TFPI2 has been identified as a tumor suppressor gene in colorectal, pancreatic, and brain cancer. In renal carcinoma, both transcript and protein levels of TFPI2 are elevated and associated with poor prognosis^42^. Thus, monitoring TFPI2 levels could therefore be valuable in assessing the progression and treatment response in cancer patients, especially those with renal carcinoma^43^ or endometrial cancer^44^.

In the second part of the study, we explored the connections between the canonical and noncanonical components of TGF-β signaling and ccRCC progression. To this means, we used the Olink panel biomarker measurements in blood from the ccRCC patients and investigated the correlation of protein expression in solid tumors from the same cohort. We mainly focused on key proteins in the TGF-β and VHL hypoxia pathways using immunoblotting as previously reported^17–19^.

For the TGF-β pathway, represented by TGFBR1-Full length receptor (FL) and TGFBR1-intracellular domain (ICD) described in Mu et al.^20^, we report here for the first time, positive associations with ADAMTS15 and FCRLB. We also found associations between the tumor promotors VEGFA, EGF, IL6, NT5E, and MSLN. The identification of a correlation between the TGF-β pathway and NT5E is of particular interest as NT5E is emerging as a crucial factor in immunoregulation in cancer biology with attempts to target NT5E in ongoing clinical studies^45–48^. We observed that overexpression of TGFBR1 and TGF-β stimulation of 786-0 ccRCC cells, which lack VHL expression, increased the expression of NT5E. Moreover, the TGFBR1 and NT5E was observed to be colocalized in the cell membrane and cytoplasm in two different ccRCC cell lines. We found negative associations between TGFBR1-FL and TGFBR1-ICD and the tumor suppressors ERBB2, SEZ6L, ICOSLG, GPNMB, FGFBP1, FASLG, TRAIL, and CTSV. Similarly, the association of TGF-β signaling components with LY9, CD48, and XPNPEP2 are reported here for the first time.

The associations with SMAD2/3 - downstream effectors in the TGF-β canonical pathway - follow the observations for TGFBR1-FL and TGFBR1-ICD, with the exception that TCL1A (TCL1 family AKT coactivator A) is negatively associated with SMAD2/3 but not associated with the upstream TGF-β receptor components TGFBR1-FL and TGFBR1-ICD. TCL1A expression is restricted to early B cells, where it acts as a coactivator of the serine threonine kinase Akt and through other interactions favoring cell survival, growth, and proliferation^49^. This association indicates that SMAD2/3 and TCL1A are connected through another pathway other than the TGF-β pathway.

The analysis of the hypoxia pathway in ccRCC tumors shows that HIF-2α was positively associated with the tumor promoters NT5E and CA9. The associations of HIF-2α and VHL are all novel observations. In summary, the PLA analysis of tumor samples support our previous finding that that the TGF-β and hypoxia pathways are interconnected^14,19^.

We used a new protein detection platform to detect biomarkers in the blood of patients with ccRCC with the aim of identifying novel biomarkers. We applied statistical modeling and reduced 91 putative biomarkers down to 7 biomarkers that can be used for a rapid test of blood samples from ccRCC patients. The study highlights the potential of applying PLA technology to identify new markers suitable for precision diagnostics. The signature panel was able to discriminate between healthy and affected individuals with high precision and high reliability. The identified proteins have all previously been reported as biomarkers for cancer, and in some cases, specifically for RCC and ccRCC. Notably, many markers are secreted proteins, and their detection in this study supports the notion that these proteins are suitable biomarkers for an analysis of blood serum.

After validation, the panel can be used for early detection, monitoring of treatments, and staging of tumors in ccRCC patients. Considering the lack of clinical symptoms at early stages of ccRCC, the usefulness for blood testing for early detection of ccRCC cannot be underestimated. It is also feasible to reiterate our study to include early stages of RCC. For this purpose, a retrospective study using the Northern Sweden biobank would be possible. Our study highlights the usefulness of technology that can extract protein signatures from small amounts of blood reposited in biobanks. Retrospective studies on such material can be invaluable for the identification of markers in future studies.

We have shown that TGF-β and hypoxia play a significant role in ccRCC tumor progression and present here novel associations between these pathways and serum biomarkers. Together, our study revealed putative novel liquid biomarkers for ccRCC, and possibly other cancer types, that can be used for development of new diagnostic tools for precision medicine. The knowledge of these biomarker interactions with TGF-β and VHL-HIF pathway signaling components in solid tumors can give a better understanding of ccRCC tumor biology, apart from augmenting diagnostic and prognostic capacity of biomarkers.

## Materials and Methods

### Patient and control participant recruitment

A total of 142 ccRCC patients from northern Sweden who underwent nephrectomy (64 men and 78 women) with a median age of 66.50 years (range 32-87 years) were included in the study. The blood and solid tumor samples were collected only after obtaining written and informed consent from the patients. All patients were surgically treated between 2000 and 2009 and samples that passed the routine quality checks were considered. The control participants who donated blood for the study were recruited, comprising 60 men and 61 women with an age range between 21-79 years and a mean age of 51.02 years. We collected surplus EDTA plasma samples from healthy blood donors, and they were made unidentifiable by removing most of the study subject information. Only age in year and gender information were kept. The use of surplus samples has been approved by the ethical board in Uppsala (01/367).

### ccRCC patient follow-up and clinicopathological parameters

The tumor size of each patient was determined by computed tomography (CT). The tumor stage distribution was determined according to the 2009 TNM classification system^24^. The Fuhrman et al.^25^ grading system was used to grade the tumors. Patient follow-up using a scheduled program was employed for survival analysis.

### Protein extraction from blood and tumors

A peripheral blood sample from the ccRCC patient cohort was collected before commencing any treatment. Similarly, blood was also collected from the healthy control group. The blood was collected in EDTA-coated tubes and centrifuged at 2000 x g for 3 hours to separate plasma. The plasma was stored at -80°C until use. Proteins from the frozen tumor tissues were extracted as previously described^17^. Briefly, a small piece of frozen tumor tissue of approximately 125 cubic millimeters was minced with a surgical knife, immediately mixed with 50 µl TRAPeze® 1X CHAPS Lysis Buffer (EMD Millipore, Billerica, MA, USA), and incubated for 30 min on ice with agitation. The mixture was then centrifuged at 14,000 rpm for 30 min, yielding a supernatant containing proteins. The total protein concentration was determined using the Thermo Scientific™ Pierce™ BCA™ Protein Assay Kit (Thermo Fisher Scientific, Waltham, MA, USA) following the manufacturer’s instructions.

### PEA of blood plasma proteins

To quantify the levels of putative oncogenic proteins in blood from ccRCC patients and healthy controls, we performed a proximity extension analysis assay (PEA) using the Olink multiplex Oncology II panel (Olink Proteomics, Uppsala, Sweden), which allows for the detection of 92 oncology-related protein biomarker candidates. The process and outcome were subjected to technical validation and quality control. The assay was carried out as previously described^50,51^. In short, 1 μl EDTA plasma sample was mixed with 3 μl incubation mix (containing 92 pairs of probes and 4 controls) in a 96-well microtiter plate. The mixture was incubated at 4°C overnight. Thereafter, 96 μl extension mix containing polymerase and PCR reagents was added. The mixture was incubated for 5 min at room temperature, 20 min at 50°C, followed by 17 cycles of DNA amplification. After completion, 2.8 μl of the amplification product was added to 7.2 μl detection mix in a new 96-well microtiter plate, from which 5 μl was transferred to a 96.96 Dynamic Array IFC (Fluidigm, South San Francisco, CA, USA), prepared and primed according to the manufacturer’s instructions. The expression analysis was performed using the BioMark™ HD real-time PCR platform (Fluidigm, South San Francisco, CA, USA). The PEA real time PCR data were exported and normalized using Olink Wizard for GenEx software. A five-parameter log-logistic function was fitted to the standard curve measurements after outliers had been removed in a procedure based on the Grubbs test^52^. The limit of detection (LOD) was defined as the protein concentration in the fitted standard curve that corresponded to the PCR cycle threshold mCtblank − 2 sCtblank, where mCtblank and sCtblank denote the mean and standard deviation threshold cycle (Ct) values for the blank, respectively. The Ct values were normally distributed, allowing use of the t-test for comparison between groups. For further statistical analysis, the Olink-defined Normalized Protein eXpression (NPX) units were used (log2 scale). Full names and UniProtKB accession numbers for all 92 proteins are listed in Supplementary Data 13.

### Power analysis to determine sample size

To ensure that the study had sufficient statistical power to distinguish between ccRCC patients and controls, power analysis was carried out using the R package MSstats^53^ using a false discovery rate of 0.05 and 92 proteins as features. The level for the power analysis of sample size was set to >0.8. The power analysis showed that 60 patients and 60 controls would be sufficient. Given this study’s sample size of 134 ccRCC cases and 111 controls, the present study has a statistical power of approximately 0.85 which is sufficient for the desired level of significance (Figure 1F).

### Statistical analyses

After collecting NPX data from the PEA experiments, principal component analysis (PCA) was performed to evaluate inter-assay variation (technical bias). Linear regression was then used to detect effects of age and gender on the same data, and the NPX values were adjusted if a significant effect was detected (p-value < 0.05). The correlation between pre- and post-adjusted NPX values for the proteins was calculated and plotted using the R software^54^ ggplot2^55^.

An initial test was performed to identify protein biomarkers (NPX data) significantly associated with either the disease (ccRCC) or control groups. The associations were estimated by a Wilcoxon signed-rank test.

The R package corrPlot^56^ was used to calculate and present several correlations within the ccRCC cohort in this study. Firstly, the correlation between the biomarkers and clinicopathological parameters -cancer grade, cancer stage, and tumor diameter-were examined. Further, the correlations between the biomarkers were analyzed. Finally, the correlations between biomarkers and protein levels obtained from solid tumor were also analyzed. All the correlation values were obtained by executing a Spearman correlation test using the corrPlot, and a *p-*value less than 0.05 was considered significant. Kaplan-Meier survival plots were plotted using SPSS (IBM) and represented cancer-specific survival in ccRCC patients.

### Multivariate analysis and Logistic Regression

To evaluate the potential of the proteins in distinguishing tumor samples from control samples, we used the top 50 most significantly altered proteins from the previous Wilcoxon signed-rank test to train a random forest (RF) model using the R package randomForest^57^. The R package ggplot2^55^ was used to visualize the importance of each protein in classifying the data. Two RF plots were produced, one ranking the proteins according to predictive value and the reduction in accuracy when proteins are excluded from the model. The more accuracy is decreased when the protein is excluded, the more important the protein is for classification. The second plot shows the Gini coefficients. The mean decrease in the Gini coefficient measures how each variable contributes to the homogeneity of the nodes and leaves in the resulting random forest, with a higher value indicating the higher importance of the variable in the model. The predictive performance of the RF model was estimated by a 10-fold cross-validated RF model using a training set (80%) and a validation set (20%). The result from the validation test was plotted as a Receiver-Operator Characteristic curve (ROC). The R package “pROC^58^ was used to plot the ROC curve and for calculations of area under the curve (AUC) and other metrics, including optimal threshold, sensitivity, specificity, positive predictive value (PPV), and negative predictive value (NPV).

In order to identify a minimal diagnostic signature protein panel for ccRCC from the top 50 most significantly altered proteins, we used an elastic-net penalized logistic regression (ENLR) model in R package glmnet^26,59^. The ENLR model combines lasso and ridge regularization penalties to simultaneously perform regularization and variable selection. The data used for the model comprise a randomly selected training set composed of 80% of samples from patients and controls, and a validation set composed of the remaining 20% of samples. The penalization proportion (α) was defined by a grid search using 10-fold cross-validation and evaluated using sensitivity, specificity, AUC, and misclassification rates, a penalty of α = 0.2 was selected for the model’s penalty proportion, as it showed the highest accuracy and lowest error rate for misclassifications. To minimize the deviance of the fitted model, the tuning parameter λ was defined as the mean value of 100 iteratively computed lambda values. Regression coefficients were determined for each protein to evaluate the contribution of each protein toward the discrimination between disease and control samples. Following the 10-fold cross-validation, a subset of proteins that showed nonzero regression coefficients in all validation passes was selected for calculation of the final regression coefficients. A final set of seven ccRCC signature proteins was defined in a procedure in which an ROC curve was generated for the protein with the highest regression coefficient and then compared to an ROC curve produced with the addition of one more protein; the protein set was considered complete when adding more proteins did not produce further significant improvement in the ROC.

### Analysis of tumor protein expression by immunoblotting (IB)

After the biomarker discovery using PEA followed by statistical analysis, we analyzed candidate biomarker expression in solid tumor samples by immunoblotting. Proteins were extracted from solid tumors as described previously^17–19^. In brief, a 125 mm piece of frozen tumor tissue was minced with a surgical knife, immediately mixed with 50 µl TRAPeze® 1X CHAPS Lysis Buffer (EMD Millipore, Billerica, MA, USA), and incubated for 30 min on ice with agitation. The mixture was then centrifuged at 20 000 x g for 30 min, yielding a supernatant containing total protein. The protein concentration was determined using the Thermo Scientific™ Pierce™ BCA™ Protein Assay Kit (Thermo Fisher Scientific, Waltham, MA, USA) following the manufacturer’s instructions. Immunoblotting was performed as previously described^17–19^. In short, Precast NuPAGE Novex (3–8%, 4–12%, 7%, 10% and 12%) gels (Life Technologies, Carlsbad, CA, USA) were used to separate 30 µg of total protein in an XCell SureLock™ Mini-Cell (Life Technologies). Trans-Blot® Turbo™ Transfer Starter System (Bio-Rad Laboratories, Hercules, CA, USA) was used to transfer the separated proteins onto a nitrocellulose membrane, preassembled in the Trans-Blot® Turbo™ Midi Nitrocellulose kit (Bio-Rad Laboratories). The membranes were incubated in Odyssey® Blocking Buffer (LI-COR Biosciences, Lincoln, NE, USA) for 60 min. The blots were then washed and incubated with diluted antibodies on a shaker at 4°C overnight. After washing, the blots were probed with the following antibodies: IRDye® 800CW goat anti-rabbit (LI-COR #926-32211, LI-COR Biosciences) or IRDye® 680RD goat anti-mouse (LI-COR #925-68070, LI-COR Biosciences). The immunoblot images were developed at 84 µm resolution in an Odyssey® CLx Infrared Imaging System (LI-COR Biosciences) The analysis of protein band densitometry was performed with Image Studio™ Software, version 3.1 (LI-COR Biosciences). The expression levels of all proteins were normalized against the expression the reference housekeeping protein, β-actin. Antibodies are listed in Supplementary Data 14. Associations between tumor protein biomarkers and components of TGF-β signaling and HIF-1α and HIF-2α and assessed using Spearman’s Rho (significant at p < 0,05). Note that the TGF-β synonyms TGFBR1-full length (FL) (Activin-like kinase 5) and TGFBR1-ICD (Intra Cellular Domain) are used in the immunoblot data and figures.

### Cell culture and *in vitro* studies

Caki-1 (VHL+/+, Sigma) and 786-O (VHL-/-, ATCC, authenticated in 2021) clear renal cell carcinoma (ccRCC) cell lines were selected based on the presence or loss of VHL expression. Caki-1 cells were grown in Dulbeccós Modified Eaglés Medium - high glucose (DMEM, Sigma-Aldrich) supplemented with 10% fetal bovine serum (FBS, Sigma-Aldrich), 1% penicillin/ streptomycin (PEST, Sigma-Aldrich), 1% L-glutamine (L-Glut, Sigma-Aldrich). The 786-O cells were grown in RPMI-1640 Medium (RPMI, Sigma-Aldrich), supplemented with 10% FBS, 1% PEST, 1% L-Glut. Media was changed every three days. Cell splitting was done with 1:5 ratio, when the cells were confluent, using Trypsin-Ethylenediaminetetraacetic acid solution (Sigma-Aldrich). Conditions for growth were 37°C and 5% CO_2_. Cells were seeded in 10-cm plates for tissue culture. When the confluency reached 60-80%, cells were transfected or directly starved depending on the experimental conditions. Transfection of 5 µg of HA-TGFBR1 or NT5E-Flag (PCMV6-NT5E-myc+Flag, Origene #RC209568) or pcDNA 3.1 was performed using FUGENE HD transfection reagent (Promega) following manufacturer instructions, for transient overexpression experiments. After 24h of transfection, cells were starved using DMEM or RPMI media, supplemented with 1% FBS, 1% PEST, 1% L-Glut for 16 h. Then, cells were treated with 10 ng/mL TGF-β1 (R&D system) for the selected time frame. Cells without treatment served as control. After the indicated time points, cells were harvested for protein extraction.

### Protein extraction

Cells were lysed using Radio Immuno Precipitation Assay buffer (RIPA) containing TRIS buffer (pH-8, 1 M), NaCl (5 M), glycerol (10%), NP4O (1%), sodium deoxycholate (0.5%), Milli-Q water. Samples were vortexed and incubated for 20 min on ice and further centrifuged at 10000xg for 10 min. Supernatant was collected and protein concentration in total protein per µL was measured using the PierceTM Bicinchoninic Acid Assay (BCA, Thermo Scientific) following manufacturer’s instructions.

### Immunoblotting of HA-TGFBR1 overexpression

A fraction of the total cell lysates was treated with sample reducing agent 10x (Thermo Fisher) and LDS sample buffer 4x (Thermo Fisher) and denatured at 95°C for 5min. From the denatured mix, 10-20 µg of protein (following maximum loading volumes) were separated on NuPageTM 7% Tris-Acetate gel (1.5mm X 10well) (Thermo Fisher) at 130V for 90min. PageRulerTM Prestained (Thermo Fisher) was used as ladder. Proteins were transferred onto a 0.2µm nitrocellulose membrane by using iBlotR2 NC Regular Stacks (Thermo Fisher) and the iBlot 2 system (Thermo Fisher) at 25V for 7min. Membranes were blocked for 1h at room temperature in TBS InterceptR Blocking Buffer (LI-COR), followed by overnight primary antibody (HA-mouse-Cell Signaling #2367, NT5E-rabbit-Cell Signaling #13160, Actin-mouse-Sigma-Aldrich #A1978) incubation at 4 °C. Membranes were washed with Tris buffered saline with Tween 20 (TBST) five times for 15 min and then incubated in secondary antibody (anti-rabbit 800CW, LI-COR #926-32211 and anti-mouse 680CW, LI-COR #926-68070) for 1 h at room temperature followed by another set of washes with TBST. Images from these membranes were obtained using an Odissey CLx (LI-COR) and Image Studio (Version 5.2, LI-COR).

### Immunoprecipitation of HA-TGFBR1 with NT5E

The total cell lysates that were obtained from the cell culture plates were incubated overnight with the primary antibody (Anti-HA, mouse, Cell Signaling), followed by a 2h incubation with protein G SepharoseTM 4 fast flow beads (Cytiva) and then beads were washed 4 times with RIPA buffer and beads were sucked dry after final wash. Once the beads were dry, they were added with LDS sample buffer and sample reducing agent and were loaded into a NuPageTM 7% Tris-Acetate gel as described above for immunoblotting. Blocking was performed with 5% bovin serum albumin (BSA) in TBST for 1h at room temperature. The membranes were incubated overnight with primary antibody (NT5E, Rabbit, Cell Signaling) at 4°C, washed with TBST five times for 15min, and incubated for 1h at room temperature with the secondary antibody (light chain specific, Cell Signaling) followed by seven washes with TBST each for 10min. Images were then obtained using Amersham Imager 600 after treatment of the membranes with Amersham ECL Prime Western Blotting Detection Reagent (Cytiva) following manufacturer’s instructions.

### Immunocytochemistry of endogenous TGBFR1 and NT5E

The Caki-1 and 786-O cell lines were grown in 6-well plates on top of glass coverslips. When the cells reached 80% confluency, they were starved for 16h followed by stimulation with 10 ng/ml TGF-β (R&D system, Minneapolis, USA) for 0min, 30min, or 3h. The medium was removed, and the cells were fixed with 4% formaldehyde for 10min at room temperature. Permeabilization was performed with 2% Triton x100 (2%) in PBS for 10min. Samples were blocked in 5% BSA in 0.2% Triton x100 in PBS. For the staining, 50µL of primary antibody (TGFBR1-rabbit-V22-1:150, NT5E-mouse-Abcam #ab257309-1:50) diluted in blocking solution were used for 1h at room temperature. Same conditions were used for the incubation with the secondary antibodies (anti-rabbit AlexaFluor555, Invitrogen #A21428-1:600, anti-mouse AlexaFluor488, Invitrogen #A32731-1:600). Mounting of cover slips on the glass slides was done using 25µL of Fluoromount containing DAPI, which stains the nuclei.

## Data availability

All data used and generated in the current work are supplied as Supplementary Data 1-14.

## Code availability

The code to replicate the work is available at Landstrom GitHub repository https://github.com/malandstromlab/Biomarkers.

## Acknowledgements

ML is funded from ALF (RV 996277 and RV 993591), Cancerforskningsfonden i Norrland/CFFN LP24-2364, UmU 982061, Swedish Cancer Society (23-2902-Pj-01-H), Swedish Research Council (2023-02370), and Erling Persson Foundation (2023-0148). CE is funded by the Knut and Alice Wallenberg Foundation under the SciLifeLab and Wallenberg National Program for Data-Driven Life Science. Initial data analysis was carried out by Genevia Technologies (Tampere, Finland).

## Ethics

The present study was approved by the institutional review board and ethical committee of Northern Sweden (Regionala Etikprövningsnämnden, Umeå University, Umeå, Sweden; approval numbers 2012-418-31M, 2018-296-32M, 2019-02579) and ethical board in Uppsala University (01/367). Written informed consent was obtained from the patients after oral information given by the staff.

## Contributions

Study concept and design: BL, MKM, and ML. Data collection: PM, RTS, RIB, and KA. Quality control of data and algorithms: PM, CE, and HH. Data analysis and interpretation: CE, PM, HH, and ML. Statistical analysis: PM and CE. Manuscript preparation: PM, CE, HH, and ML. Manuscript editing: PM, CE, HH, RTS, KA, AL, BL, MKM, and ML. All authors have read and approved the submitted article version.

